# PRISM-M: A Recurrent Framework for the Formation of Stable Internal Neural Models

**DOI:** 10.64898/2026.09.02.748818

**Authors:** Eliezer Masliah

## Abstract

How transient neural representations become integrated and stable enough to function as internal neural models remains incompletely understood. Grounded in efficient coding, Bayesian and predictive frameworks, recurrent and attractor dynamics, neural state-space models, and systems neuroscience, the Principle of Representation Integration for Stable Models (PRISM) proposes five operations: extraction, compression, integration, stabilization, and prediction/action. Here we developed PRISM-M, a minimal nine-equation recurrent dynamical realization with an explicit contraction condition (0 < *J* < 1), to examine whether these operations can generate persistent, context-sensitive, and prospectively informative model states. Seven simulation analyses showed persistent but revisable trajectories and a 0.209 context-dependent shift in the event-period model state. A 60 × 60 parameter sweep identified 719 rigid, 2,344 adaptive, and 537 high-gain trajectories within this structurally contractive parameter space, and the same regimes were recovered across 1,000 randomized environments. Perturbations of extraction, integration, and stabilization altered model trajectories, whereas the scalar compression perturbation had a small effect. The full PRISM state predicted the next model state more accurately than the model-state-only baseline (RMSE 0.0186 versus 0.0218), while performing similarly to an unconstrained ARX model. In the BART dataset, spatial fMRI states were distinguishable in 155 participants (69.7% accuracy; 33.3% chance), and inflation-related activity was modestly associated with pumping behavior in the 99 participants with matched behavioral data (*β* = 0.198, *P* = 0.040). PRISM-M provides a constrained, testable framework centered on four operational signatures of model-like organization structured representation, contextual integration, persistence or reconstructability, and prospective relevance.

**Author Summary:** The brain continually receives information from the outside world, the body, and its own ongoing activity, yet useful behavior requires more than simply detecting these signals. Neural information must be selected, organized, combined with context, maintained over time, and used to guide what happens next. We developed PRISM-M, a simple recurrent mathematical implementation of the Principle of Representation Integration for Stable Models (PRISM), to examine how these steps may work together within an explicit model. PRISM-M treats extraction, compression, integration, stabilization, and prediction/action as five interacting operations and tests whether they can produce internal states that persist over time while remaining able to change. Across seven simulations, the model generated context-sensitive and revisable states, remained mathematically stable across broad parameter ranges, and showed predictable effects when individual operations were altered. We also examined an independent human fMRI dataset from a sequential risk-taking task. The available data supported structured, condition-sensitive neural patterns and a modest relationship with behavior. They also indicated that temporally resolved recordings will be needed to test the full recurrent model directly. Together, these results provide a quantitative and testable framework for studying how neural representations may become stable internal neural models.

## Introduction

The nervous system is continuously exposed to signals arising from the external environment [1, 2], the internal state of the organism [3], and its own ongoing activity across distributed cortical and subcortical networks [4–6]. Many of these signals are transient, yet perception, memory, decision-making, regulation, and behavior depend on neural states that extend beyond the moment in which a signal is encountered. The same external event can acquire different significance depending on physiological state, prior experience, current goals, or recent history. Neural processing therefore requires relevant structure to be selected, represented in a usable form, related to other available information, and maintained sufficiently long to influence what the nervous system does next.

Several major theories address components of this problem. For instance, efficient-coding and information-bottleneck approaches describe how relevant information can be preserved while redundancy is reduced [1, 2, 7, 8]. Bayesian and predictive-processing theories emphasize interactions between incoming evidence, prior knowledge, and expectations [9, 10], while active-inference and control frameworks extend these relationships to action and regulation [11, 12]. Recurrent and attractor models provide mechanisms for persistence or reinstatement [13, 14], and memory-related studies show how previous experience shapes new information [15, 16]. Dynamical-systems and neural-manifold approaches emphasize evolving population states [17, 18], whereas interoceptive, allostatic, and network frameworks emphasize bodily state and distributed implementation [3–5].

What remains less explicitly formulated is the intermediate transformation between the availability of structured signals and the formation of an internal neural model that can subsequently be used. Current theories begin with different questions, including how information is encoded efficiently, how priors influence inference, how prediction errors drive updating, how neural states are maintained, how actions are selected, or how population activity evolves through state space. Here the Principle of Representation Integration for Stable Models (PRISM) addresses a complementary organizational question: what functional transformations allow signals arising from the world, body, memory, and ongoing neural activity to become a coherent, sufficiently stable, and usable internal model? PRISM places these contributions within a common signal-to-model process at an intermediate level between neural coding and downstream inference, prediction, regulation, and behavior.

PRISM organizes this transformation into five interacting functional operations: extraction, compression, integration, stabilization, and prediction/action. Extraction selects relevant structure from the information available to the nervous system. Compression transforms selected information into a more economical representation while preserving distinctions needed for subsequent processing. Integration relates that representation to organismal state, context, prior knowledge, memory, goals, value, and other ongoing information. Stabilization provides sufficient continuity for the resulting organization to persist or be reconstructed while remaining revisable. Finally, an established model can influence subsequent inference, prediction, simulation, regulation, or action. These operations are functionally distinguishable but dynamically coupled and may overlap, recur, and operate at different spatial and temporal scales.

PRISM is intended as a systems-neuroscience framework whose operations emerge through interactions among distributed neural systems, including visual, somatomotor, dorsal attention, ventral attention/salience-related, limbic, frontoparietal control, and default mode networks [4, 5]. Attention and salience systems contribute to selection and processing priority [19, 20]; limbic and interoceptive systems convey affective, motivational, and bodily relevance [3]; frontoparietal systems support flexible goal-dependent control [21]; and default mode and hippocampal-cortical systems contribute memory-based context and internally generated states [15, 16]. These cortical networks interact with thalamic, basal ganglia, cerebellar, brainstem, autonomic, and other subcortical systems [3, 5]. PRISM therefore proposes distributed, overlapping implementation rather than one network for each operation.

This organization also emphasizes that model formation is neither entirely outside-in nor entirely inside-out [6, 9, 12]. External sensory signals, interoceptive and proprioceptive information, action-related feedback, and intrinsic neural activity provide evidence from which models can be formed and revised [3, 11, 15]. Established models, in turn, can influence what information is sampled, how ambiguous signals are interpreted, and which actions are considered. PRISM therefore places internal models within a recurrent brain-body-environment cycle. Supporting the role of this recurrent organization, recent work showing coordinated transitions between sensory encoding and internally oriented memory retrieval across brain-wide activity [6].

In this context, an important distinction arises between a neural representation and an internal neural model [11, 15, 22]. A neural state may reliably encode a stimulus, feature, relationship, or condition without necessarily functioning as a model. In PRISM, model-like organization develops as represented information becomes integrated with other relevant states, acquires sufficient stability to persist or be reinstantiated, and becomes capable of constraining subsequent processing. An internal neural model can therefore be considered a biologically instantiated organization of neural states and their relationships or transformations that can be elicited or reinstantiated by relevant signals, is sufficiently stable to persist or be reconstructed, and can be operated upon to support inference, prediction, simulation, regulation, or action. This shifts the observational question from whether information can be decoded from neural activity to whether a neural organization has the integration, continuity, and functional consequences expected of a model.

PRISM thus connects neural representation, contextual integration, distributed network coordination, temporal stabilization, and prospective model use as distinguishable but interacting requirements of model formation. This organization also creates experimentally separable questions. A system may encode a signal but integrate it incompletely, form an integrated state that is not maintained, or establish a stable model that is used differently for prediction or action.

A conceptual framework alone cannot establish whether these operations can function together within a coherent dynamical system or how persistence and updating are balanced over time. A useful internal model should withstand transient fluctuations while remaining sufficiently flexible to incorporate meaningful change. Prospective influence from an established state should also be distinguishable from effects attributable simply to current input or persistence of the preceding state. These questions motivated PRISM-M, a minimal mathematical implementation that translates the biological logic of PRISM into an explicit dynamical framework.

Here we formulate PRISM-M as an ordered recurrent system in which extracted information is compressed, integrated with contextual and prior-state information, and incorporated into a stabilized model state with prospective influence. Seven simulation analyses examine persistence and revisability, context dependence, stability–plasticity regimes, operation-specific perturbations, prospective prediction, and robustness across heterogeneous conditions. An independent Balloon Analogue Risk Task (BART) behavioral and fMRI dataset is then used as an empirical boundary test of structured spatial neural states and their relationship to behavior. The study asks whether this signal-to-model architecture can generate persistent, context-sensitive, revisable, and prospectively informative states with experimentally distinguishable consequences.

## 2. Mathematical formulation of PRISM-M

PRISM-M was developed to determine whether the biological logic proposed by PRISM could be expressed as a coherent recurrent dynamical system and once expressed mathematically, whether its individual operations could be examined independently. The formulation was intentionally kept minimal. It is not intended as a mathematical reconstruction of the brain or as a set of equations derived from fixed neurophysiological constants. Rather, the equations provide an explicit representation of the functional relationships proposed by PRISM so that their interactions, stability, perturbations, and prospective consequences can be examined quantitatively. The current Version 4 formulation represents the locked nine-equation realization used for the synthetic analyses.

The mathematical formulation progresses through three levels. Equation 1 places PRISM-M within the general class of recurrent dynamical systems. Equation 2 specializes this general architecture according to the ordered functional organization proposed by PRISM. Equations 3–8 provide the minimal scalar realization used to make extraction, compression, integration, stabilization, and prospective model use executable. Finally, Equation 9 is derived from the preceding equations and provides an analytic measure of effective recurrence and contraction. Thus, the nine equations serve different roles within the formulation. Some define the overall architecture, others implement the proposed biological operations, and the final equation describes a mathematical property of the resulting system.

Before introducing the individual equations, the basic signal-to-model transformation can be summarized schematically as:

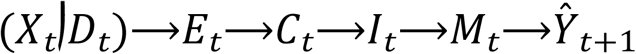

Here, *X_t_* represents structured information available to the system, *D_t_* represents disturbance or competing information, *O_t_* represents organismal or contextual state, and *M_t−1_* represents the preceding model state. The intermediate variables *E_t_*, *C_t_*, and *I_t_*represent successive stages of model construction, whereas *M_t_*represents the stabilized model state and *Y_t+1_* a subsequent neural, cognitive, regulatory, or behavioral consequence.

### 2.1. General recurrent dynamical framework

At the most general level, PRISM-M can be placed within a recurrent dynamical framework in which the next state of a system depends on its current state, incoming information, contextual influences, and variability:

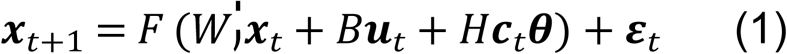

In Equation 1, *x_t_* represents the current state of the system, *u_t_* incoming information, and *c_t_* contextual influences. *W*, *B*, and *H* determine how these components contribute to the next state, *F* represents the transformation of these inputs, *θ* contains model parameters, and *ε_t_*represents unmodeled variability.

Equation 1 is conceptually important because it captures a basic biological premise of PRISM. A neural response reflects both the signal arriving at a given moment and the state already present in the system, together with information about the organism and its context. Equation 1 therefore defines the broader dynamical architecture within which PRISM-M operates, while the simulations are generated using the more specific equations that follow (3-9).

### 2.2. PRISM-specific ordered architecture

PRISM-M next specializes this general recurrent framework according to the functional sequence proposed by PRISM:

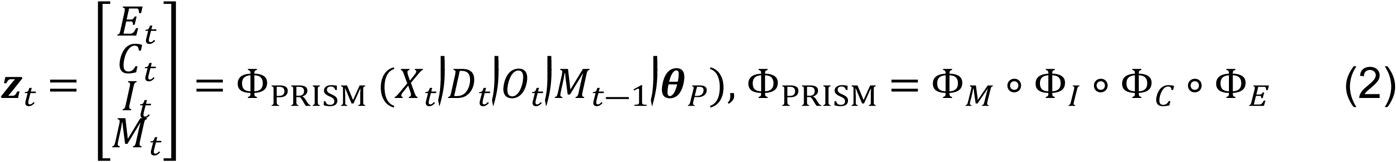

Equation 2 converts the general recurrent architecture (Equaton 1) into a PRISM-specific sequence. In expanded form, extraction maps the structured signal and disturbance to *E_t_*; compression maps *E_t_* to *C_t_*; integration combines *C_t_*, *O_t_*, and *M_t−1_* to generate *I_t_*; and stabilization combines *I_t_* with *M_t−1_* to generate *M_t_*. These relationships clarify the ordered composition without adding further core equations.

The sequence is functional rather than anatomical. Extraction, compression, integration, and stabilization are not proposed to occur in four separate brain regions or as isolated serial events. Their biological implementation is expected to involve overlapping and recurrent interactions among distributed neural systems. Equation 2 simply specifies the order of transformations required for the minimal mathematical realization. In the current scalar model, *E_t_*, *C_t_*, and *I_t_* are instantaneous transformations, whereas temporal recurrence is carried primarily through *M_t_*and its dependence on *M_t−1_*.

### 2.3. Extraction

Here, represents the structured signal and competing, distracting, or otherwise less relevant information. The intact model gives greater weight to the structured signal while retaining a smaller contribution from disturbance:

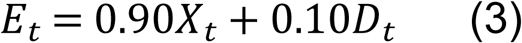

Here, *X_t_* represents the structured signal and *D_t_* competing, distracting, or otherwise less relevant information. The intact model therefore gives greater weight to the structured signal while allowing some contribution from disturbance.

This is a deliberately simple representation of a broad biological process. In neural systems, extraction could involve receptive-field selectivity, attentional selection, gain modulation, salience detection, signal-to-noise enhancement, or combinations of these mechanisms. Equation 3 captures the general functional requirement that relevant structure be preferentially retained for subsequent processing.

The values of 0.90 and 0.10 were selected as illustrative parameters that provide a high-fidelity extraction condition while preserving a measurable contribution from disturbance. Other weightings that maintain preferential selection of the structured signal would represent the same functional operation. For the degraded extraction condition, the weights are shifted to 0.70 and 0.30, increasing the influence of disturbance and providing a controlled reduction in extraction fidelity.

### 2.4. Representational compression

The extracted signal is next transformed into a finite-precision representation:

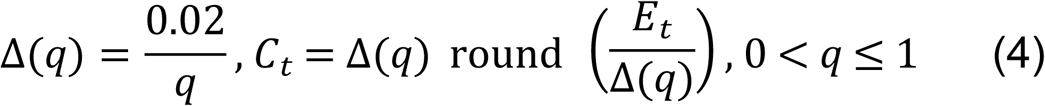

The parameter *q* controls representational precision, while the constant 0.02 sets the reference spacing between representational levels. At *q*=1, the quantization step is 0.02, and progressively smaller values of q increase this spacing and produce a coarser representation. The value 0.02 was selected as an illustrative reference scale that provides relatively fine encoding at high *q* while allowing representational precision to vary over the range examined in the simulations.

Equation 4 is best understood as a minimal model of lossy scalar encoding. It allows information to be represented with different levels of precision while maintaining the overall scale of the signal. It is not intended to imply that biological neural compression occurs through literal numerical quantization. In the brain, compression may instead involve sparse or efficient population coding, dimensionality reduction, abstraction, manifold organization, or other transformations that preserve useful structure while reducing representational complexity.

This formulation separates representational precision from signal amplitude. In Version 4, changing *q* primarily alters the spacing between representational levels, allowing the effects of compression fidelity to be examined independently of systematic changes in signal magnitude.

### 2.5. Contextual integration

PRISM proposes that a representation acquires model-like relevance in part by being related to the current state of the organism and to information already present in the system. This is represented as:

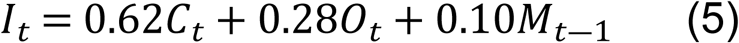

Equation 5 combines three sources of information. *C_t_* carries the currently represented evidence, *O_t_* represents organismal or contextual state, and *M_t−1_*carries information from the preceding model state. The coefficients 0.62, 0.28, and 0.10 were selected as illustrative weights that give the greatest contribution to the current representation, a substantial contribution to context, and a smaller contribution from the preceding model state. Their sum of one keeps the integrated state on a comparable scale. Other weighting schemes could represent the same functional operation while changing the relative influence of current evidence, context, and prior model state.

The implication is that he same external representation does not necessarily generate the same internal state. Its significance can depend on physiological condition, motivational state, task context, memory, previous experience, goals, or the model already active in the system. *O_t_* is therefore an aggregate contextual variable rather than a proposed activity measure of one brain region or network.

The inclusion of *M_t−1_* allows prior model organization to shape the interpretation of newly represented information. This provides the recurrent contribution to integration and connects contextual integration with the stability analysis developed below.

### 2.6. Persistence and input gain

After integration, the system balances persistence of previously established organization with responsiveness to newly integrated information. PRISM-M represents these two requirements through a persistence transformation, *p(s),* and an input-gain transformation, *u(η).* The first determines how strongly the preceding model state is retained, while the second determines how strongly new integrated information contributes to the evolving model state.

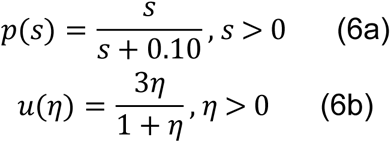

Together, Equations 6a and 6b constitute Equation 6 in the nine-equation PRISM-M formulation.

The first transformation converts the raw stabilization parameter *s* into a persistence term *p*(*s*). For positive values of *s*, *p*(*s*) remains between 0 and 1. Increasing *s* therefore increases persistence while keeping the effective recurrent contribution bounded.

The second transformation, *u*(*η*), determines how strongly newly integrated information can influence the evolving model. Unlike *p*(*s*), *u*(*η*) is an input gain and is not restricted to values below one. Persistence and responsiveness to new information can therefore be adjusted independently.

This separation could be is biologically useful as a model may be highly persistent but relatively insensitive to new evidence, or it may retain prior organization while responding strongly to meaningful changes in integrated input. PRISM-M uses these two terms to represent the balance between stability and plasticity without treating them as opposite ends of a single parameter.

### 2.7. Stabilized model-state evolution

The preceding components are combined in the central recurrent update equation:

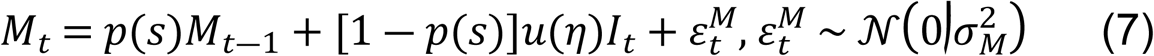

Equation 7 describes how the internal model state evolves over time. The first term preserves a proportion of the preceding model state, whereas the second introduces newly integrated information. The noise term represents variability not explicitly modeled; in the stochastic simulations it is additive, state-independent, zero-mean noise with finite variance.

Equation 7 therefore describes a recurrent updating process in which part of the preceding model state is retained while newly integrated information contributes to the evolving state. An important point is that Equation 7 is not a simple convex average between old and new information. Because *u*(*η*) is an independently adjustable input gain and may exceed one, the coefficients multiplying the previous and new states do not generally sum to one.

Biologically, Equation 7 represents the central stability-plasticity problem addressed by PRISM. If persistence dominates, an established model may resist meaningful updating. If newly integrated information has excessive influence, the state may respond strongly to each change in input. Between these conditions lies a range in which the model can maintain continuity while remaining revisable. The simulations below examine whether these different regimes emerge from the equations rather than imposing them as separate model types.

### 2.8. Prospective model deployment

A central distinction between a representation and an internal model is that an established model should be capable of influencing what occurs subsequently. PRISM-M represents this prospective relationship as:

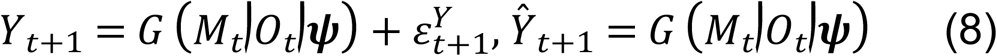

Here, *M_t_* is the stabilized model state, *O_t_*the organismal or contextual state, and *Y_t+1_* a subsequent measurable outcome. Depending on the experimental setting, that outcome could be a future neural state, prediction, regulatory response, decision, behavior, or action. *G* represents the mapping through which the established model influences that outcome, and *ψ* contains the corresponding parameters.

Equation 8 is intentionally general because the biological implementation of model use is likely to differ substantially across neural systems and behaviors. A sensorimotor model, for example, may influence an upcoming movement, whereas an interoceptive model may influence autonomic regulation and an episodic model may contribute to simulation or future-oriented behavior.

The purpose of Equation 8 in the present formulation is limited but critical as it establishes that the model state is not simply an endpoint of processing but it can have prospective consequences. The intermediate computations involved in model reinstantiation, simulation, evaluation, selection, action, feedback, and revision are not decomposed here.

In Simulation 5, this prospective implication is evaluated using predefined predictive models and run-wise cross-validation rather than by assuming a specific biological form for *G*.

### 2.9. Effective recurrence and contraction

Because Equation 5 feeds the preceding model state *M_t−1_* back into integration, and Equation 7 then uses the integrated state to construct the next model state, the effective recurrent influence can be derived directly. Substituting Equation 5 into Equation 7 and isolating the dependence on *M_t−1_*gives the effective recurrent coefficient:

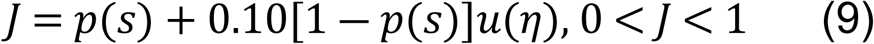

The coefficient *J* represents the total influence of the preceding model state on the updated state. It combines the direct persistence of *M_t_−1* in Equation 7 with its additional recurrent contribution through integration in Equation 5. Larger values of *J* therefore correspond to stronger carryover of the preceding state, whereas smaller values correspond to more rapid attenuation across successive updates.

Equation 9 is derived from the recurrent structure specified by Equations 5 and 7 and provides a mathematical property of the resulting system. For all admissible positive values of *s* and *η* in the present scalar realization, *J* remains between 0 and 1 by construction. The recurrence is therefore contractive throughout the admissible parameter space, so differences in the preceding model state progressively decrease over successive updates. This structural condition provides mathematical stability while allowing a range of trajectory patterns.

This distinction is important for interpreting the rigid, adaptive, and high-gain regimes examined in the simulations. A high-gain trajectory can produce large responses or greater volatility while remaining mathematically contractive. “High-gain” therefore describes the responsiveness of the model trajectory within a stable recurrent system.

The present stability result applies to the scalar realization under the assumptions used here, including fixed parameters within a run, bounded inputs, and additive finite-variance state-independent noise. The scalar model can be extended to a multidimensional state *M_t_*, with recurrent coupling represented by a matrix or state-dependent Jacobian. In a linear vector extension, asymptotic stability would require the spectral radius *ρ(J)<1*, while nonlinear extensions would use corresponding local or global stability conditions. This provides a direct route from the present scalar realization to distributed population-state models.

### 2.10. From representation to an operational internal neural model

The equations provide a mathematical counterpart to the distinction introduced earlier between a representation and an internal neural model. The unnumbered schema at the beginning of this section summarizes the progression from available signals through extraction, compression, integration, stabilization, and prospective model use.

The first three transformed variables represent different stages of model construction. *E_t_* contains preferentially extracted evidence. *C_t_* represents that evidence at a defined level of fidelity. *I_t_* relates the representation to contextual and prior-state information. None of these stages alone is assumed to constitute an internal neural model.

In the present realization, *M_t_* is the first variable explicitly assigned the role of a stabilized model state because it combines integrated current information with continuity from the preceding state. Equation 8 then adds a prospective criterion by asking whether that state can constrain what occurs next.

This leads to four operational properties that can guide expeimental identification of model-like neural organization: i) structured representation, ii) integration with relevant contextual or prior information, iii) persistence or reconstructability beyond the immediate input, and iv) prospective relevance for subsequent neural, regulatory, cognitive, or behavioral outcomes. These properties are intended as experimentally useful signatures rather than as a claim that *M_t_* corresponds to a single neural population or anatomical location.

An internal neural model might be distributed across multiple regions, expressed as an evolving neural trajectory, reconstructed through recurrent interactions, or partly latent in synaptic and network organization between episodes of active expression.

### 2.11. Relationship between the PRISM architecture and the tested realization

It is useful to distinguish three levels of the present framework. First, PRISM defines the biological process architecture through the proposed operations of extraction, compression, integration, stabilization, and prediction/action. Second, PRISM-M translates that architecture into a recurrent mathematical framework that specifies how these operations relate dynamically. Third, the present Version 4 formulation provides one specific minimal scalar realization of PRISM-M, using the equations and parameter values tested in the simulations.

The coefficients in Equations 3–7 are illustrative parameters chosen to illustrate the functional relationships in the present scalar realization. Finite-precision quantization provides one simple implementation of compression, and Equation 7 provides one explicit mechanism for stabilization. The same functional requirements could be expressed through alternative equations, higher-dimensional population states, different timescales, nonlinear transformations, or distributed network interactions.

The simulations that follow in the next section use this nine-equation realization to determine whether these properties produce persistent yet revisable states, context dependence, distinguishable stability-plasticity regimes, operation-sensitive perturbation effects, and prospective information.

For fixed parameter values, the scalar PRISM-M state equation can be written in a first-order autoregressive form with transformed exogenous inputs and is therefore similar to ARX(1) (Ljung, 1999). However, PRISM-M builds on this established recurrent form by organizing the input-to-state transformation into interpretable operations of extraction, compression, integration, and stabilization, followed by prospective model use. The main difference is that PRISM-M places a general recurrent dynamical structure within a biological framework for internal neural model formation. While ARX(1) provides a general description of how previous states and external inputs determine subsequent states, PRISM-M specifies candidate neural transformations through which incoming information is selected, organized, integrated with organismal context and prior state, stabilized, and subsequently used.

## 3. Computational and empirical methods

PRISM-M was evaluated through seven simulation analyses followed by an independent database analysis of behavioral and fMRI data from the Balloon Analogue Risk Task (BART) set (Wei and Qin, 2025). The simulations examined complementary aspects of the signal-to-model process, including persistence and revisability, context dependence, stability–plasticity behavior, operation-specific perturbations, prospective prediction, and robustness across heterogeneous conditions.

All simulations used the Version 4 realization described in Section 2, representing the final prespecified implementation used throughout the study. Once this version was established, the equations, parameter definitions, perturbations, and simulation procedures were held fixed. Equations 1 and 2 define the general recurrent architecture and PRISM-specific organization, whereas Equations 3–9 provide the executable realization used in the simulations. The same implementation and random seed 20260731 were used to support reproducibility. The BART analysis was treated separately as a real-world based test of selected observable features relevant to PRISM-M rather than as a direct fit of the nine equations.

### 3.1. Numerical implementation and common model measures

The simulated model followed the PRISM sequence defined in Section 2. Here Equation 3 represented extraction of structured information from competing input, Equation 4 compression through finite-precision encoding, Equation 5 integration of the compressed representation with context and the preceding model state, and Equations 6 and 7 stabilization through persistence, input gain, and model-state evolution. Equation 8 represented the prospective prediction/action component examined in Simulation 5. Equation 9 was not a separate PRISM operation; it provided the corresponding mathematical measure of effective recurrent stability. Unless otherwise indicated, the basic simulated environment consisted of 140 time steps divided into baseline, event, and post-event periods. Structured evidence *X_t_*changed across these periods, while disturbance *D_t_* was generated as a temporally correlated competing signal. The default realization used *η* = 0.60, *s* = 1.20, and *q* = 0.85, with additive model-state noise having a standard deviation of 0.01. The default event began at time 40 and ended before time 85.

Parameters were held constant within each run. Inputs were bounded, and stochastic simulations used additive, state-independent, zero-mean Gaussian noise with finite variance, consistent with the stability assumptions in Section 2. Baseline activity was calculated from the 20 time points preceding event onset. Event responses for the integrated state *I_t_* and model state *M_t_* were defined as the difference between event-period and baseline means. The adaptation ratio was the absolute response of *M_t_* divided by the absolute response of *I_t_*, providing a measure of how strongly the stabilized model followed changes in the integrated representation. Volatility was calculated from successive changes in *M_t_* during the event and normalized to model-response magnitude. Root-mean-square error (RMSE) and next-state error were retained as additional descriptive measures. Three operational dynamical patterns were defined before examining the parameter-sweep results. A trajectory was classified as rigid when its adaptation ratio was below 0.50. It was classified as high-gain when the adaptation ratio exceeded 1.25 or normalized volatility exceeded 0.40. Trajectories falling between these conditions were classified as adaptive. These thresholds are operational criteria for the present realization rather than proposed biological boundaries.

For each simulation run, the effective recurrent coefficient from Equation 9 was calculated to document where the run lay within the structurally stable range. This allowed the observed behavior of the model to be considered separately from the magnitude of effective recurrence: a trajectory could respond strongly or show greater variability while the underlying recurrent dynamics remained stable.

### 3.2. Simulation analysis design

#### 3.2.1. Simulation 1: Basic model dynamics

Simulation 1 examined whether the complete signal-to-model sequence could generate a persistent but revisable internal state. The default environment and model parameters were used, and trajectories of external evidence, integrated representation *I_t_*, and stabilized model state *M_t_*were followed across baseline, event, and post-event periods. The analysis focused on the temporal relationship between *I_t_* and *M_t_* and on persistence after the initiating input changed. This simulation used Equations 3–7 to generate the successive stages from extraction through stabilized model-state formation. Equation 9 was calculated as a numerical check of the structural contraction condition. The simulation was intended as a general test of the model dynamics and was not designed to reproduce a particular sensory system, neural circuit, or behavioral task.

#### 3.2.2. Simulation 2: Context-dependent model formation

Simulation 2 asked whether the same represented evidence could produce different model states when contextual state changed. The evidence sequence, disturbance, timing, model parameters, and noise were held constant, while *O_t_* was assigned three levels: low (*O* = 0.25), intermediate (*O* = 0.55), and high (*O* = 0.85). In this synthetic analysis, *O_t_* was an abstract contextual variable representing, in simplified form, influences such as internal state, prior information, or task context. Separate model-state trajectories were generated for each condition. This analysis primarily examined Equation 5, in which contextual information enters the integrated state, and followed the consequences through Equations 6 and 7 into *M_t_*. Because the other inputs were identical, differences among trajectories reflected the contribution of *O_t_* rather than changes in incoming evidence.

#### 3.2.3. Simulation 3: Stability-plasticity parameter sweep

Simulation 3 addressed how persistence and responsiveness to new information interact in determining model-state behavior. A 60 × 60 parameter sweep varied the update/input-gain parameter *η* from 0.02 to 1.40 and stabilization strength *s* from 0.25 to 2.50, yielding 3,600 parameter combinations. Compression quality was held at *q* = 0.85, environmental input was fixed, and stochastic model noise was removed so that differences across parameter space reflected deterministic model dynamics. This simulation primarily examined Equations 6 and 7. Equation 6 determines persistence *p*(*s*) and input gain *u*(*η*), whereas Equation 7 determines how these two components combine to update *M_t_*. For every parameter combination, the adaptation ratio and normalized volatility were calculated, and the resulting trajectory was assigned to the rigid, adaptive, or high-gain category defined above. Because Equation 9 guarantees 0 < *J* < 1 for all admissible positive values of *s* and *η*, the sweep examined these trajectory patterns within a recurrently stable parameter space. The purpose of the parameter sweep was to determine whether the same recurrent architecture could express different balances between persistence and updating as its parameters changed.

#### 3.2.4. Simulation 4: Effects of selective changes in PRISM operations

Simulation 4 tested what happens to the model trajectory when individual PRISM operations are selectively altered. Four prespecified perturbations were compared with the intact Version 4 realization. Degraded extraction altered Equation 3 by changing the weighting of structured signal and disturbance from 0.90/0.10 to 0.70/0.30. Poor compression altered Equation 4 by reducing *q* to one half of its intact value, with a lower bound of 0.10, thereby creating a coarser finite-precision representation. Importantly, this manipulation changed representational fidelity without introducing the amplitude reduction that had been present in the earlier Version 3 formulation. No integration altered Equation 5 by setting *I_t_* = *C_t_*, thereby removing the explicit contributions of organismal context and the preceding model state. Weak stabilization reduced *s* by 50%, altering the persistence term in Equation 6 and its effect on model-state evolution in Equation 7.

The intact and perturbed realizations used the same evidence, context, event timing, and noise sequence. The principal perturbation measure was RMSE between the altered *M_t_* trajectory and the corresponding intact trajectory; changes in event response and other dynamical measures were also retained. The perturbations were selected to produce interpretable changes in different components and were not calibrated to equivalent perturbation magnitude. Differences in RMSE therefore describe sensitivity of the present realization rather than an intrinsic ranking of the PRISM operations.

#### 3.2.5. Simulation 5: Prospective model-state prediction

Simulation 5 investigated whether an established PRISM-M state carried useful information about the next model state. This analysis addressed the prospective component represented generally by Equation 8, using states produced by Equations 3–7. A separate prediction dataset was generated from 150 independent simulated runs, each containing 120 time steps. Event onset and termination, evidence levels, contextual levels, signal variability, and model parameters varied between runs. For these simulations, *η* ranged from 0.35 to 0.80, *s* from 0.70 to 1.80, *q* from 0.70 to 0.95, and model-noise standard deviation from 0.005 to 0.025. The final time point from each run was excluded because there was no subsequent state to predict, leaving 17,850 prospective observations.

The primary target was *M_t+1_*. Rather than assuming that a particular PRISM variable should be sufficient, several predictor sets were compared. This included evidence only (*X_t_*, *D_t_*), extraction only (*E_t_*), compression only (*C_t_*), integration only (*I_t_*), model state only (*M_t_*), a context-enriched representation (*C_t_*, *O_t_*), and the full PRISM state (*E_t_*, *C_t_*, *I_t_*, *M_t_*, *O_t_*). Three reduced reference representations were also included: a free ARX-like model using (*M_t_*, *C_t_*, *O_t_*), a no-quantizer model using (*M_t_*, *E_t_*, *O_t_*), and a no-recurrence model using (*C_t_*, *O_t_*). A conventional linear-Gaussian ARX/Kalman reference was evaluated separately.

The comparison was designed to address several related questions. The *M_t_*-only model tested how much prospective information could be explained by persistence of the current state alone. Evidence-only and no-recurrence models asked whether the next state could be predicted primarily from current input without an established recurrent model state. The full PRISM predictor tested whether combining current representation, integration, context, and the stabilized state added information beyond persistence alone. An unconstrained autoregressive model with exogenous inputs (ARX) and a linear-Gaussian Kalman model [23, 24] were included as standard comparison models to determine whether prospective prediction could also be achieved by more general recurrent approaches. A separate no-quantizer model tested whether the finite-precision compression step contributed additional predictive information.

Prediction was evaluated using five-fold GroupKFold cross-validation implemented in scikit-learn [25]. All observations from a given simulated run were kept within the same fold, preventing neighboring time points from the same trajectory from appearing in both training and test data. Performance was quantified using RMSE and *R*^2^. The main comparison was between the full PRISM predictor and the *M_t_*-only model. RMSE was therefore also calculated separately for each of the 150 independent runs. Paired run-level differences were summarized using 5,000 bootstrap resamples and evaluated with Wilcoxon signed-rank tests. The same run-wise procedure was used for comparisons with the reduced reference models.

Within PRISM-M, prospective prediction was treated as a candidate functional signature of model-like organization rather than as a definition of an internal neural model. An internal neural model would be expected to influence subsequent neural or behavioral states, but prospective information alone does not establish that a neural state has the broader properties of an internal model.

#### 3.2.6. Simulation 6: Randomized robustness

Simulation 6 evaluated whether the same dynamical patterns remained evident when both the environment and the model parameters varied. One thousand independent environments were generated. Event onset varied between time points 30 and 49 and event termination between 75 and 99. Baseline, event, and post-event levels of evidence and context were randomized together with the amount of signal and distractor variability. Across runs, *η* ranged from 0.02 to 1.40, *q* from 0.20 to 1.00, *s* from 0.25 to 2.50, and model-noise standard deviation from 0.01 to 0.12.

Simulation 6 extended the stability–plasticity analysis of Simulation 3 to heterogeneous conditions. Instead of varying persistence and input gain within a fixed environment, signal, context, event timing, model parameters, and noise varied across 1,000 runs. The same adaptation, volatility, and trajectory criteria were applied, and Equation 9 was calculated to document effective recurrence within the structurally stable range. This tested whether the dynamical patterns identified in the systematic sweep remained evident across a broader range of simulated conditions.

#### 3.2.7. Simulation 7: Randomized perturbation analysis

Simulation 7 repeated the same operation-specific perturbations across heterogeneous environments. The four changes used in Simulation 4 were applied to each of the 1,000 randomized environments generated for Simulation 6. Within an individual run, the intact and perturbed models shared the same evidence, context, parameter values, event timing, and stochastic noise.

This paired design allowed the effect of changing a particular operation to be separated from differences among environments. Equation 3 was perturbed for extraction, Equation 4 for compression, Equation 5 for integration, and Equations 6–7 for stabilization. RMSE relative to the corresponding intact trajectory and changes in event response were calculated for every perturbation and run.

### 3.3. Empirical boundary test using BART behavioral and fMRI data

#### 3.3.1. Dataset, analytical scope and ethics statement

The expeimental analysis used the BART, a laboratory paradigm developed to examine risk-taking behavior [26]. BART was selected because it combines condition-specific neural measurements with behavior during sequential decisions, providing an external test of distinguishable neural states and behavioral relevance. The analysis used the Behavioral and ROI-level fMRI Data from the Balloon Analogue Risk Task dataset [27].

The behavioral component included 99 participant files containing 8,717 balloon trials. The fMRI dataset contained 200 parcel files for each of three conditions intact, inflation, and explosion with 155 participants represented in every parcel file. All 99 participants with behavioral data were represented in the fMRI dataset and formed the matched sample for neural–behavioral analyses. Inspection of the uploaded fMRI files established the empirical scope of the analysis. The 600 files were parcel-specific matrices in which rows represented participants and columns represented spatial features within each parcel, rather than participant-specific BOLD time series. Across the 200 parcels, the files contained 151,349 spatial features, with feature counts varying by parcel.

The available BART data therefore support condition-sensitive spatial and between-participant analyses, but not tests of preceding-state persistence, event-by-event updating, temporal recurrence, hemodynamic lag, or Kalman state transitions. Accordingly, the PRISM-M equations were not fitted directly to BART data. The analysis asked whether the supplied neural measurements contained structured, condition-sensitive spatial states and whether static summaries of those states were related to behavior. It therefore served as an external boundary test of representation and behavioral relevance rather than a test of Equations 3–9.

The experimental component of this study consisted of secondary analysis of previously collected, publicly available, de-identified behavioral and fMRI data from the Balloon Analogue Risk Task dataset reported by Wei and Qin [27] and deposited in Mendeley Data (Version 1; DOI: 10.17632/xjn4n9cvxs.1). The present study did not recruit participants, interact with participants, collect new human-subject data, or access directly identifiable private information. Ethical approval and informed-consent procedures governing the original collection of the BART data were the responsibility of the original investigators. The present analyses were limited to the publicly available de-identified dataset.

#### 3.3.2. Behavioral reconstruction and spatial neural representation

The behavioral files were combined after participant identification. Participant-level measures included trial count, mean pumps, adjusted mean pumps, explosion rate, reward, mean trial duration, and total task length. Trial-level bookkeeping relationships were checked before proceeding with neural–behavioral analyses. Sequential behavioral analyses were used to characterize the task data but were kept separate from the simulated PRISM-M dynamics.

For each parcel and experimental condition, three participant-level neural summaries were calculated across the available spatial features: mean signal, signal standard deviation, and mean absolute signal. Global measures were calculated across all 151,349 spatial features using feature-count weighting. Paired condition contrasts were examined across the 155 participants represented in the full fMRI dataset. Benjamini– Hochberg false-discovery-rate correction was applied separately within each metric and condition-contrast family across the 200 parcels [28].

Principal-component analysis was used to characterize the covariance structure of the complete 200-parcel spatial representations. These analyses were intended to determine whether the supplied neural maps could be represented in a more compact spatial form. They were not interpreted as direct evidence for the compression operation represented by Equation 4, because spatial covariance and biological compression are not equivalent.

Condition separability was tested using the 200 parcel means. Five-fold GroupKFold cross-validation was used, with all three condition maps from the same participant assigned to the same fold. This analysis therefore used the full 155-participant fMRI sample and asked whether intact, inflation, and explosion conditions could be distinguished from their distributed spatial patterns.

#### 3.3.3. Neural-behavioral associations

Associations between neural measures and behavior were examined in the 99 participants with both behavioral and fMRI data. The complete feature screen included three condition values and three condition contrasts for each of the three neural summary measures across 200 parcels, together with 18 global summaries, yielding 3,618 neural features per behavioral outcome. False-discovery-rate correction was applied across the complete set of features within each outcome using the Benjamini– Hochberg procedure [28].

Because several behavioral measures are strongly influenced by the overall structure and duration of the task, a sensitivity analysis adjusted both the neural variables and behavioral outcomes for trial count and total task time before repeating the association analysis. This provided a more conservative assessment of whether neural–behavioral relationships remained after accounting for these broad task-related differences.

A focused analysis examined global mean neural activity during balloon inflation in relation to mean pumping behavior. The primary relationship was evaluated using a linear model with HC3 heteroskedasticity-consistent standard errors [29]. Additional analyses examined the stability of the relationship across alternative covariates, transformations, winsorization, exclusion of influential observations, permutation testing, bootstrap resampling, leave-one-out analyses, and repeated cross-validation. The final locked robustness analysis used 10,000 bootstrap resamples. Quadratic, threshold/hinge, and natural cubic spline models were examined as secondary alternatives to determine whether the relationship was better described by a simple nonlinear form.

Secondary behavioral outcomes included adjusted mean pumps, explosion rate, mean reward, and mean duration. Total reward was treated as a derived sensitivity variable because it is mathematically related to trial count and mean reward rather than providing a fully independent behavioral endpoint.

#### 3.3.4. Incremental behavioral prediction

A separate analysis asked whether static neural information improved prediction of participant behavior beyond information already available from the general structure of the task. The earlier static analysis was first audited to determine how its features and preprocessing steps had been constructed. The corrected analysis then used the complete 200-parcel dataset, with all scaling, dimensionality reduction, and model selection performed inside the training folds to prevent information from the test data from entering model construction.

Prediction used 10-fold cross-validation repeated five times. Candidate neural representations included global neural summaries, 600 parcel means, 1,800 parcel summaries, and 1,800 condition contrasts. Task-structure-only models were compared with neural-only and combined task-plus-neural models using the same behavioral outcomes. Ridge-model tuning was performed within each training set. Participant-level bootstrap intervals and paired sign-flip tests were used to assess incremental changes in prediction error. These machine-learning analyses were implemented using scikit-learn [25].

These analyses addressed the behavioral relevance of static participant-level neural states. The recurrent temporal sequence central to PRISM-M—including contextual recurrence, stabilization, prospective deployment, and state-to-state dynamics— requires temporally resolved data and was outside the scope of the available BART files.

### 3.4. Reproducibility and statistical analysis

All seven simulations were generated from the same Version 4 implementation rather than reconstructed separately for individual analyses. The reproducibility package retained the executable source, model specification, parameter ranges, perturbation definitions, random-number sequence, locked seed 20260731, output tables, and SHA-256 manifest. Where simulations required paired comparisons, the intact and comparison models used identical environmental inputs and noise sequences.

RMSE was used to quantify differences between intact and perturbed model-state trajectories. Prospective prediction was evaluated using RMSE and *R*^2^, with cross-validation grouped by independent simulation run. Run-level paired comparisons used Wilcoxon signed-rank tests together with bootstrap confidence intervals. Mathematical stability followed analytically from Equation 9, and the effective recurrent coefficient was also calculated for each parameter combination or randomized realization as a numerical check.

The BART analysis used a separate locked seed, 20260802. Analyses were performed in Python 3.13.5 using NumPy 2.3.5, pandas 2.2.3, SciPy 1.17.0, statsmodels 0.14.6, and scikit-learn 1.8.0. Scikit-learn was cited according to its standard publication reference [25]. Source scripts, intermediate and final outputs, figures, file manifests, and the analysis audit were retained in the dataset reproducibility package. Unless otherwise indicated, statistical tests were two-sided. Benjamini–Hochberg procedures were used to control the false-discovery rate for multiple spatial comparisons [30]. Cross-validation was grouped according to the independent unit of analysis simulation run for the prospective simulations and participant for the BART analyses to reduce information leakage between training and test observations.

Source scripts, intermediate and final outputs, figure-source data, file manifests, and the analysis audit are included in the publicly available PRISM-M Version 4 reproducibility package deposited in Zenodo (DOI: 10.5281/zenodo.22070209).

### 3.5. Use of Artificial Intelligence Tools

OpenAI ChatGPT was used as an assistive tool during preparation of this study, including support for organization and revision of manuscript text and for the development, review, and documentation of computational analyses and code. All mathematical formulations, model assumptions, computational procedures, statistical analyses, figures, numerical results, references, scientific interpretations, and conclusions were independently reviewed and verified by the author. AI-generated output was not accepted without author review, and the author takes full responsibility for the accuracy and integrity of the work and the final manuscript.

## 4. Results

### 4.1. PRISM-M generates persistent and context-sensitive model states

The first two simulations established the basic behavior of model-state formation. In Simulation 1, a transient change in the input signal generated a model state that persisted while remaining responsive to subsequent change (Figure 2A, B). The integrated state *I_t_* responded rapidly to the event, whereas the stabilized model state *M_t_* changed more gradually and retained the influence of preceding states. The adaptation ratio was 0.917, placing the trajectory within the predefined adaptive range (Figure 2D). This pattern suggests that stabilization can provide temporal continuity while preserving responsiveness to new information. Then Simulation 2 extended this behavior by examining the contribution of context. With input signal, disturbance, timing, model parameters, and noise held constant, changing *O_t_* shifted the model-state trajectory across the three contextual conditions (Figure 2C). The difference between the low- and high-context conditions was 0.209 in mean event-period *M_t_* (Figure 2D). Because the input signal was unchanged, this shift reflected the contextual contribution to integration and its subsequent influence on the stabilized model state.

**Figure 1.**
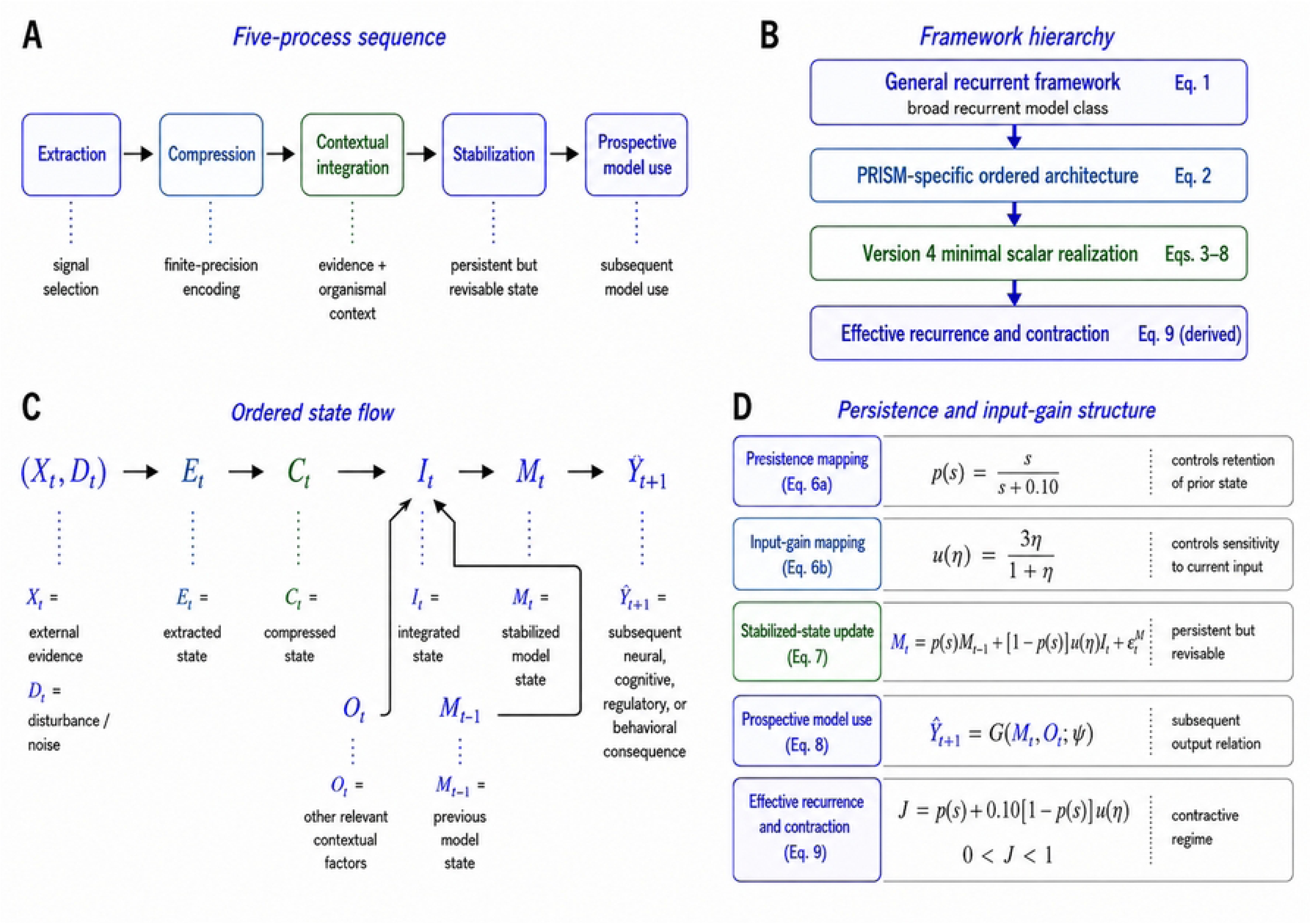
PRISM-M architecture and equation hierarchy. (A) Five-process sequence linking extraction, compression, contextual integration, stabilization, and prospective prediction/action. (B) Relationship between the general recurrent framework and the constrained PRISM-M realization. (C) Ordered state flow from structured input and disturbance through extracted (*E_t_*), compressed (*C_t_*), integrated (*I_t_*), and stabilized model (*M_t_*) states to a prospective output. (D) Schematic relationship among persistence, input gain, stabilized-state updating, and effective recurrence. The panel is conceptual; the complete locked mathematical definitions are given in Equations 1–9 in the text.

**Figure 2.**
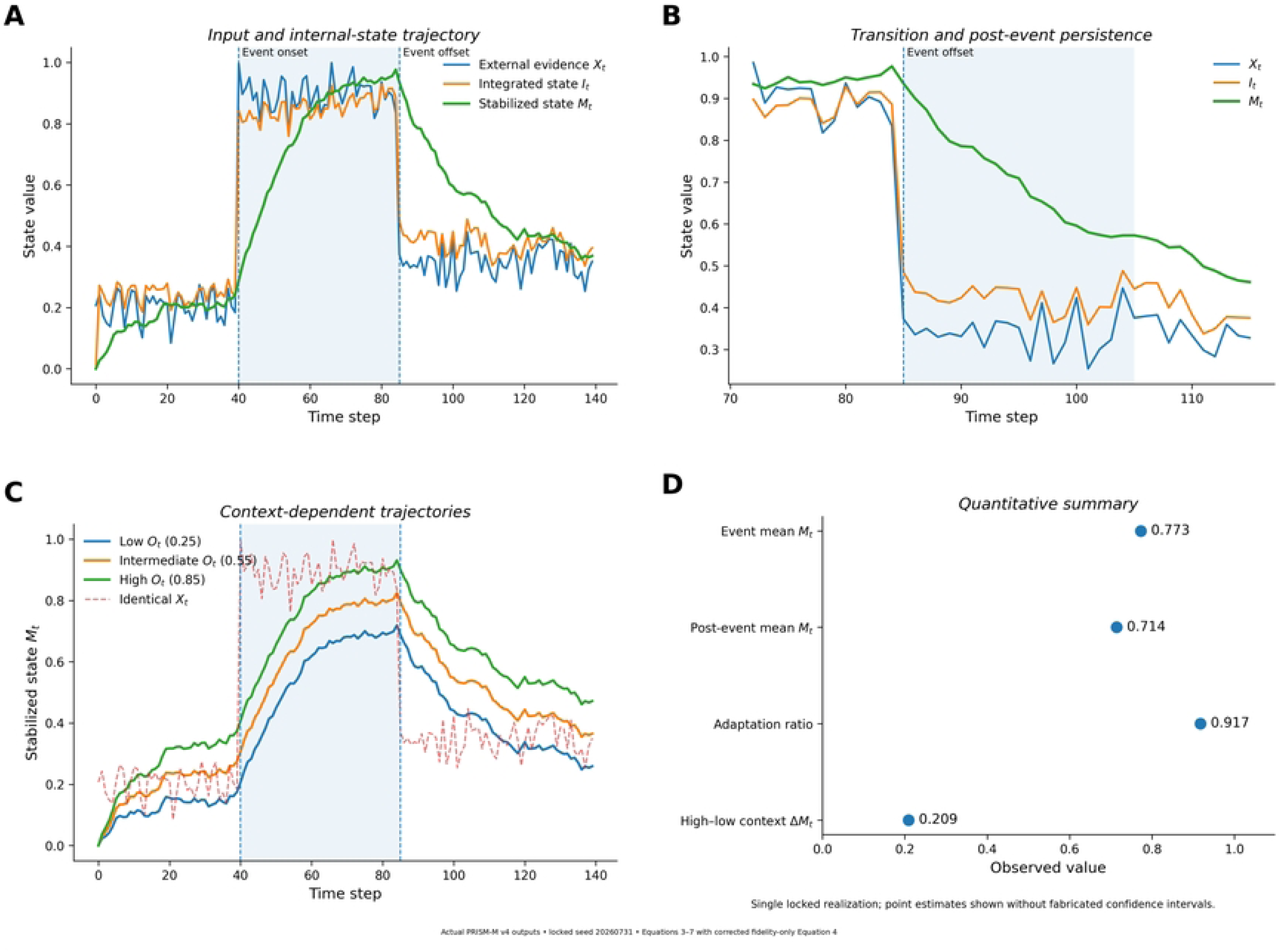
Basic dynamics and contextual modulation. (A) Simulation 1 trajectories for external evidence *X_t_*, integrated state *I_t_*, and stabilized model state *M_t_*across baseline, event, and post-event periods. (B) Expanded transition around event offset, illustrating the slower decay and persistence of *M_t_*relative to *X_t_* and *I_t_*. (C) Simulation 2 model-state trajectories for low (*O* = 0.25), intermediate (*O* = 0.55), and high (*O* = 0.85) context under an identical evidence sequence. (D) Quantitative summary of event-period *M_t_*, post-event *M_t_*, adaptation ratio (0.917), and the high–low contextual difference in event-period *M_t_* (0.209).

### 4.2. Stability–plasticity behavior emerges across parameter combinations

The next step examined how stability–plasticity patterns varied across model space. Simulation 3 evaluated how persistence and input gain shaped model-state behavior (Figure 3A, B). Across the 60 × 60 parameter sweep, the 3,600 combinations separated into three operational patterns: 719 rigid, 2,344 adaptive, and 537 high-gain trajectories. The adaptive region occupied the largest portion of parameter space, while more persistent and more responsive regimes emerged as the balance between stabilization and updating changed. Because Equation 9 guarantees 0 < *J* < 1 across the admissible positive parameter range, the sweep examined rigid, adaptive, and high-gain behavior within a recurrently stable space (Figure 3A). Trajectories with stronger responsiveness or greater variability, including high-gain trajectories, therefore remained mathematically stable. Representative trajectories illustrate the corresponding differences in model-state evolution (Figure 3B).

**Figure 3.**
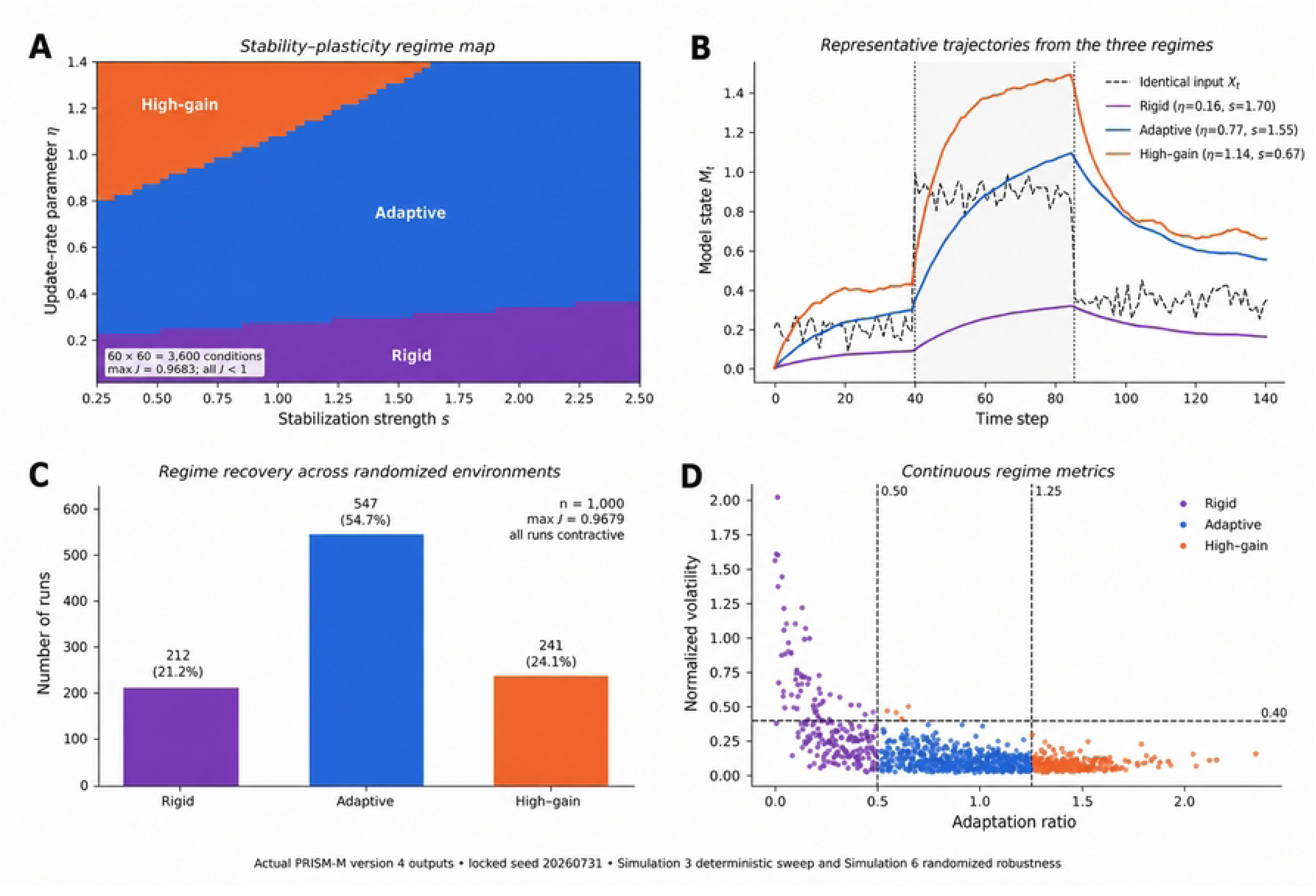
Stability–plasticity regimes and robustness. (A) Simulation 3 stability– plasticity regime map across the 60 × 60 parameter sweep (3,600 combinations), identifying rigid, adaptive, and high-gain regions as stabilization strength *s* and input sensitivity *η* vary. All combinations satisfied the recurrent stability condition in Equation 9. (B) Representative trajectories from the three operational regimes under a common input. (C) Simulation 6 regime recovery across 1,000 randomized environments: 212 rigid, 547 adaptive, and 241 high-gain runs; all remained contractive. (D) Continuous adaptation-ratio and normalized-volatility measures underlying the operational classifications.

These results suggest that the same recurrent architecture can express different balances between persistence and updating as persistence and input-gain parameters change.

### 4.3. Selective changes in PRISM operations alter model-state trajectories

The contribution of individual PRISM operations was examined by selectively altering each component and comparing the resulting trajectory with the intact model. In Simulation 4, weak stabilization produced an RMSE of 0.0854, degraded extraction 0.0842, and removal of contextual and recurrent integration 0.0781. The finite-precision compression perturbation produced a smaller change, with an RMSE of 0.00260 (Figure 4A). Because compression was defined as a change in representational precision rather than signal amplitude, the smaller effect describes the sensitivity of this scalar implementation to reduced compression fidelity. The perturbations were not calibrated to equivalent magnitude, so differences in RMSE are interpreted as properties of the tested realization rather than an intrinsic hierarchy among PRISM operations.

**Figure 4.**
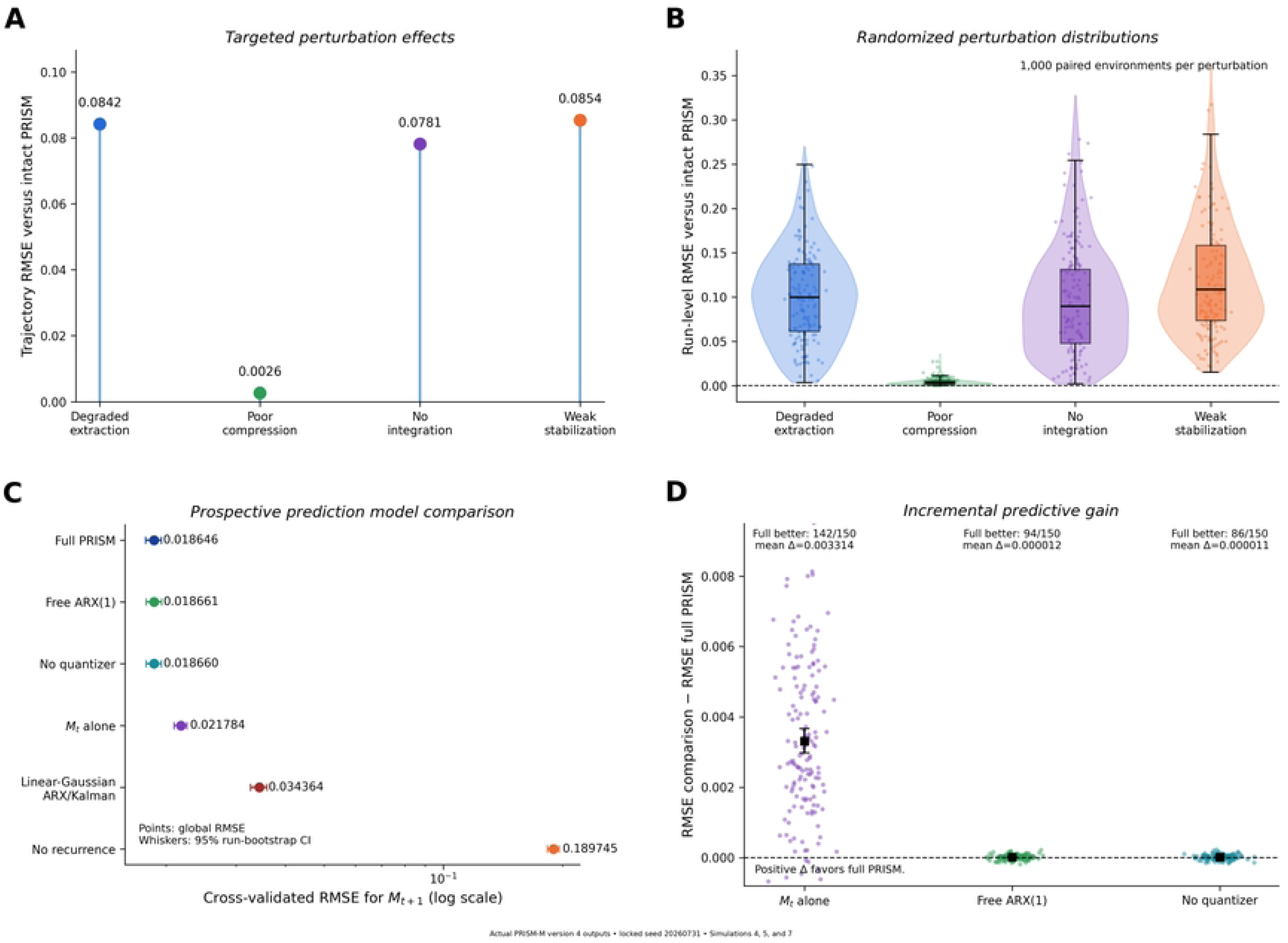
Operation-specific perturbations and prospective prediction. (A) Simulation 4 trajectory RMSE relative to intact PRISM-M after degraded extraction (0.0842), reduced-fidelity compression (0.0026), removal of integration (0.0781), or weak stabilization (0.0854). (B) Simulation 7 distributions of run-level RMSE across 1,000 paired randomized environments for the same perturbations. (C) Simulation 5 cross-validated RMSE for prediction of *M_t+1_* using the full PRISM state and comparison models. The full PRISM model achieved RMSE 0.018646, compared with 0.021784 for *M_t_* alone, 0.018661 for free ARX(1), 0.018660 for the no-quantizer reference, 0.034364 for the linear-Gaussian ARX/Kalman reference, and 0.189745 for the no-recurrence reference. (D) Run-level incremental predictive gain relative to full PRISM; the full model outperformed *M_t_* alone in 142 of 150 runs, with mean RMSE improvement 0.00331.

### 4.4. The recurrent model state carries prospective information

The prospective analysis tested whether the established PRISM-M state carried information forward in time. In Simulation 5, the full PRISM predictor achieved an RMSE of 0.018646 and *R*^2^ = 0.995384, compared with RMSE 0.021784 and *R*^2^ = 0.993700 for the *M_t_*-only model (Figure 4C). Because both predictors operated near the predictive ceiling, the absolute improvement over *M_t_*alone was small. The improvement occurred in 142 of 150 independent simulation runs, with a mean run-level RMSE reduction of 0.00331. The paired comparison yielded *P* = 1.50 × 10⁻²⁵ (Wilcoxon signed-rank), reflecting the consistency of this small run-level difference across the 150 paired simulations (Figure 4D).

The reduced models helped place this result in context. Removing recurrence increased RMSE to 0.189745 and reduced *R*^2^ to 0.522020. The linear-Gaussian Kalman reference performed better than the no-recurrence model but remained less accurate than the full PRISM representation (RMSE 0.034364; *R*^2^ = 0.984322) (Figure 4C). The unconstrained autoregressive model with exogenous inputs (ARX) and the no-quantizer reference performed very similarly to the full PRISM model, with RMSE values of 0.018661 and 0.018660, respectively (Figure 4C). Their paired run-level differences from the full model were correspondingly close to zero (Figure 4D). This convergence suggests that the prospective information captured by PRISM-M is also accessible to more general recurrent formulations, while PRISM-M provides an interpretable organization of the contributing states.

These findings suggest that prospective influence may serve as a candidate functional signature of model-like organization, as one component of the broader properties expected of an internal neural model.

### 4.5. Model behavior remains robust across heterogeneous simulated conditions

Simulation 6 extended the stability–plasticity analysis across 1,000 randomized environments in which signal, context, event timing, model parameters, and noise varied. The same three dynamical patterns remained evident: 212 rigid, 547 adaptive, and 241 high-gain runs (Figure 3C). As expected from the structural contraction condition in Equation 9, all 1,000 simulations remained within recurrently stable dynamics. Adaptive trajectories remained the largest group, while rigid and high-gain trajectories were also represented. Continuous adaptation and volatility measures further show that the operational categories arise within a broader dynamical landscape (Figure 3D).

Moreover, Simulation 7 then extended the operation-specific perturbations across the same 1,000 heterogeneous environments. Weak stabilization produced a mean RMSE of 0.1198 relative to the corresponding intact model, degraded extraction 0.1017, removal of integration 0.0971, and reduced-fidelity compression 0.00455 (Figure 4B). The pattern observed in Simulation 4 (Figure 4A) was also evident across the randomized environments in Simulation 7 (Figure 4B). Changes in extraction, integration, and stabilization continued to shift model-state trajectories, whereas the compression manipulation produced a smaller effect. Variation in effect magnitude across environments suggests that sensitivity to individual operations depends in part on the surrounding simulated conditions.

### 4.6. BART identifies structured neural states and a modest relationship with behavior

The BART analysis provided an independent empirical test of selected features relevant to PRISM-M. Because the available fMRI data consisted of participant-level spatial features rather than temporally resolved BOLD trajectories, the analysis focused on condition-sensitive spatial organization, neural–behavioral relationships, and the predictive value of static neural measures (Figure 5).

**Figure 5.**
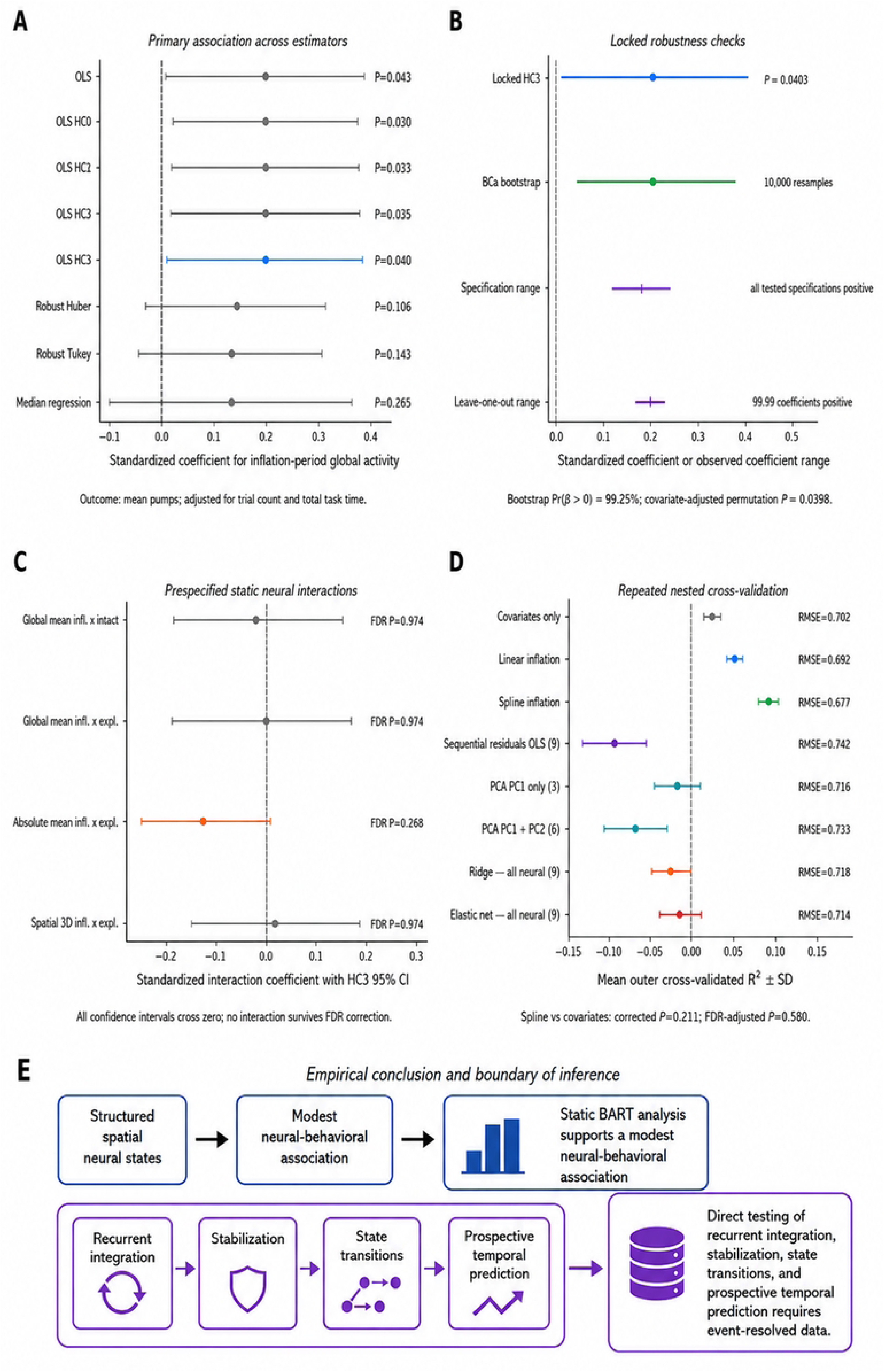
Final BART reanalysis in the 99 participants with matched behavioral and fMRI data. (A) Standardized association between inflation-period global neural activity and mean pumping behavior across alternative estimators; the prespecified HC3 estimate was *β* = 0.198 (*P* = 0.040; 95% CI 0.009–0.388). (B) Locked robustness analyses, including 10,000-resample BCa bootstrap, specification checks, and leave-one-out analysis. (C) Prespecified static neural interaction tests; confidence intervals crossed zero and no interaction survived false-discovery-rate correction. (D) Repeated nested cross-validation of task-structure and neural prediction models, showing no reliable corrected incremental contribution from static neural summaries. (E) Empirical conclusion and boundary of inference: the static BART analysis supports a modest neural–behavioral association, while recurrent integration, stabilization, state transitions, and prospective temporal prediction require event-resolved data. Condition classification reported in the text used the complete 155-participant fMRI sample and is not displayed in this figure.

Using the complete 155-participant fMRI sample, distributed activity across 200 parcels distinguished intact, inflation, and explosion conditions with 69.7% classification accuracy, compared with a three-class chance level of 33.3%. The spatial maps also retained substantial shared organization, with intact and explosion patterns showing a mean within-participant correlation of *r* = 0.890. Principal-component analysis similarly indicated compact spatial covariance structure. These findings support structured, condition-sensitive spatial representation while leaving temporal PRISM-M operations unresolved. Neural–behavioral analyses were restricted to the 99 participants with matched behavioral and fMRI data. Across the broader feature screen and prespecified static interaction analyses, no consistent corrected multivariable pattern emerged; the prespecified interaction terms remained nonsignificant after false-discovery-rate correction (Figure 5C). A focused analysis identified a modest positive association between global mean inflation-related activity and mean pumping behavior (*β* = 0.198, HC3 *P* = 0.040; 95% CI 0.009–0.388) (Figure 5A).

The direction of this association remained stable across sensitivity analyses. The bootstrap confidence interval was 0.044–0.365, 99.2% of bootstrap estimates were positive, the permutation test gave *P* = 0.040, and the coefficient remained positive in all 99 leave-one-out analyses (Figure 5B). These results support a modest relationship between inflation-related activity and pumping behavior while favoring a cautious interpretation of effect size. Repeated cross-validation showed limited incremental predictive value for static neural summaries. Adding neural representations to task-structure information did not provide a corrected improvement in prediction of mean pumps, adjusted mean pumps, explosion rate, mean reward, or mean duration (Figure 5D).

The BART findings therefore provide an initial empirical link to PRISM-M and clarify the scope of what can be evaluated with the present dataset (Figure 5E). Structured spatial states and a modest neural–behavioral association are compatible with selected features relevant to model-like neural organization. Direct evaluation of recurrent integration, stabilization, reinstantiation, and prospective state transitions will require temporally resolved neural measurements.

## 5. Discussion

The present study showed that the PRISM operations of extraction, compression, integration, stabilization, and subsequent model use can be expressed within a coherent recurrent mathematical framework and can operate together to generate states with properties relevant to internal neural models (Figure 6). The modeling analyses identified five main features of the PRISM-M architecture: (1) transient changes in input generated model states that persisted while remaining revisable; (2) contextual information shifted the resulting model state even when the incoming signal was held constant; (3) variation in persistence and input gain produced rigid, adaptive, and high-gain patterns within recurrently stable dynamics; (4) selective perturbation of individual PRISM operations produced distinguishable changes in model-state trajectories; and (5) the established PRISM-M state carried prospective information about the subsequent state. The BART analysis provided an initial extension, identifying structured spatial neural states and a modest association with behavior. Together, these findings support the internal coherence and testability of the PRISM-M architecture and suggest specific directions for further computational and experimental development.

**Figure 6.**
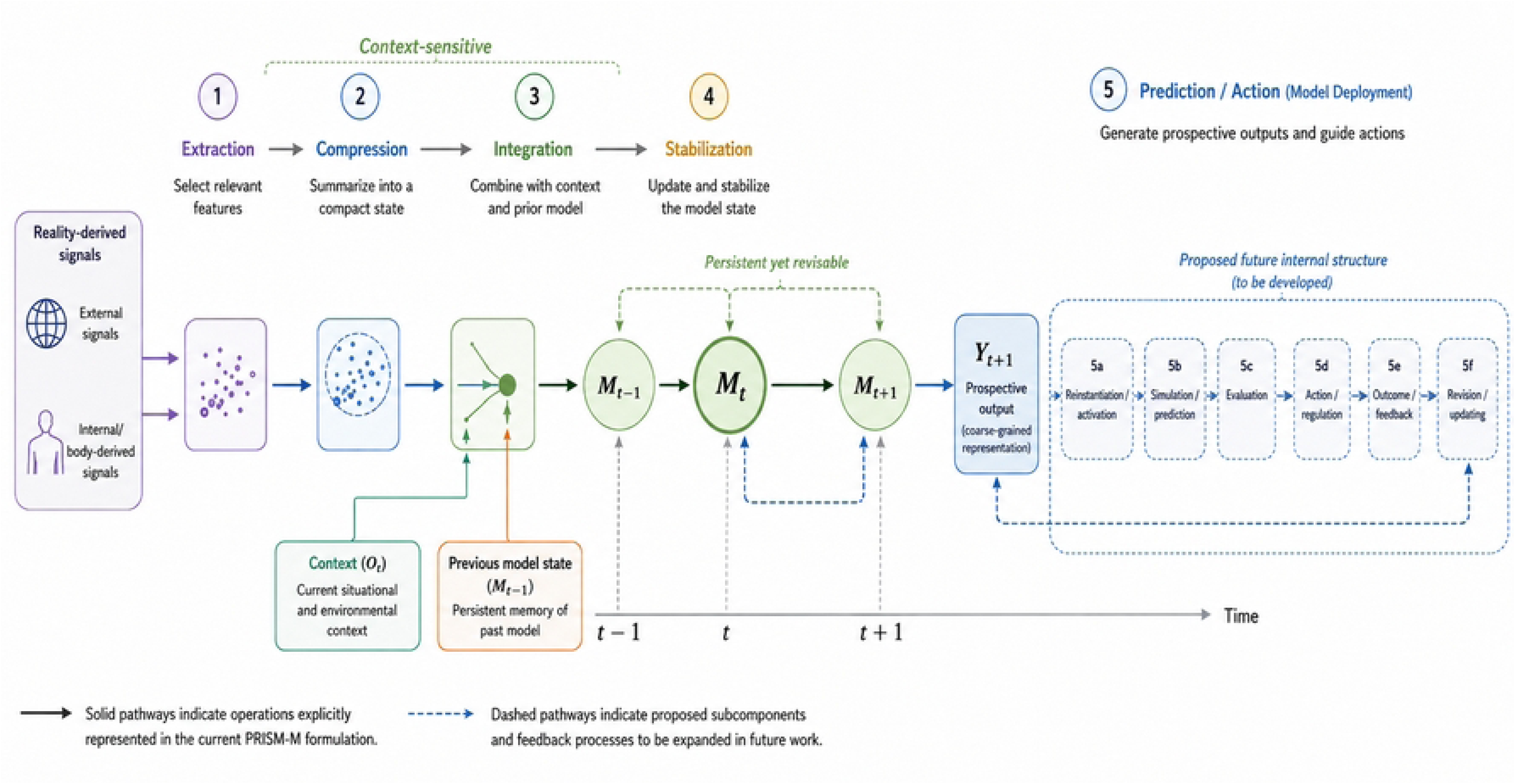
Dynamic interpretation of PRISM-M model formation and deployment. Reality-derived external and internal/body-derived signals enter extraction and compression before contextual and prior-state information contributes to integration and stabilization. The model state evolves recurrently from *M_t−1_* to *M_t_* and *M_t+1_*, illustrating persistence with continued revisability. Operation 5, prediction/action, is represented in the current PRISM-M formulation by a coarse-grained prospective output *Y_t+1_*. Dashed pathways outline proposed subcomponents for subsequent development, including model reinstantiation or activation, simulation/prediction, evaluation, action/regulation, outcome/feedback, and revision/updating. Solid pathways denote operations explicitly represented in the current formulation; dashed pathways denote proposed internal structure and feedback relationships.

The persistence observed in Simulation 1 is consistent with previous studies showing that recurrent dynamics can maintain information beyond the duration of an initiating signal while preserving the capacity to update. Attractor and integrator networks provide established examples of persistent activity, accumulation of noisy information, error correction, and longer-timescale dynamics [31]. Population-level approaches have shown that persistence can be expressed through trajectories, low-dimensional manifolds, or recurrently generated states rather than a single fixed activity pattern [32, 33]. PRISM-M treats this requirement at a functional level, with stabilization allowing an integrated state to become sufficiently persistent or reconstructable to participate in subsequent processing while remaining compatible with multiple biological implementations [32, 34]. The evolving sequence from *M_t−1_* to *M_t_* and *M_t+1_* emphasizes this dynamic rather than static interpretation (Figure 6).

Another relevant finding was that the persistent model state remained sensitive to contextual information, shifting even when the incoming signal was held constant. Changing *O_t_* modulated the model state while the input signal and other conditions were kept constant. This is consistent with experimental studies showing that context can shape neural computation; in primate prefrontal cortex, the same sensory dimensions can be selected and integrated differently according to behavioral context through recurrent population dynamics [35]. More generally, neural representations are shaped by task demands, prior information, behavioral state, and downstream use [36, 37]. In the present model, *O_t_* represents context broadly and combines several possible sources of contextual information. Future multidimensional formulations could separate factors such as interoceptive state, memory-related information, task context, and neuromodulatory influences to better examine how each contributes to model formation.

The parameter analyses also addressed the balance between persistence and updating, a central requirement for a model that must remain stable while still responding to new information. Across both the systematic parameter sweep and the randomized environments, variation in persistence and input gain produced rigid, adaptive, and high-gain trajectories while the recurrent dynamics remained stable. This relationship connects PRISM-M with the longstanding stability–plasticity problem, in which useful information must be retained while new information remains able to modify the system [38, 39]. Related robustness–flexibility trade-offs are also central to recurrent and attractor network theory [31]. In PRISM-M, Equations 6 and 7 provide a compact separation of persistence and input gain, while Equation 9 places the admissible positive parameter range within a contractive recurrent regime. The systematic and randomized analyses then examined how rigid, adaptive, and high-gain behavior can arise within these stable dynamics. These results indicate that strong responsiveness can coexist with mathematical stability. Here the terms rigid, adaptive, and high-gain therefore describe different patterns of model behavior within the parameter space examined here and are not intended to represent distinct biological categories. In higher-dimensional formulations, these patterns may instead form continuous transitions or include metastable states, multiple attractors, oscillatory dynamics, or transitions between neural manifolds [32].

The perturbation analyses provided a complementary test of the architecture. Selective changes in extraction, integration, and stabilization produced reproducible shifts in the model-state trajectory, and the same general pattern was recovered across heterogeneous environments. This supports a functional decomposition in which individual operations can be manipulated and compared mathematically. The decomposition is not intended to imply separate anatomical modules. Neural computations commonly emerge from distributed population activity in which encoding, transformation, integration, and downstream readout overlap across regions and timescales [40, 41]. The distinction between functional organization and biological mechanism is important when interpreting a model at this level [42]. PRISM-M provides experimentally addressable functional relationships; cellular, synaptic, and circuit mechanisms remain to be established for particular biological implementations.

The current scalar model likely underrepresents the biological complexity of compression. Reducing finite-precision fidelity produced only a small change in downstream trajectories. This result characterizes the tested quantization mechanism rather than the broader role of neural compression. Efficient-coding theories emphasize redundancy reduction and adaptation to input statistics, population coding transforms information into forms suitable for downstream computation, and information-theoretic approaches ask how relevant information can be retained while other variation is reduced [8, 40, 43]. A multidimensional PRISM-M formulation could examine compression through dimensionality, redundancy, representational geometry, and preservation of task-relevant or predictive information. Neural-manifold approaches provide a natural framework for such an extension [32].

The prospective analysis showed that the established PRISM-M state contained information about the subsequent model state. The full PRISM state predicted *M_t+1_* somewhat more accurately than *M_t_*alone, although both models operated near the predictive ceiling. The small improvement was highly consistent across paired runs, while removal of recurrence produced a much larger reduction in predictive performance. This pattern supports prospective prediction as a self-consistency check on information carried forward by the model state. This is consistent with internal neural model concepts in motor control and control theory, in which internal states contribute to prediction, state estimation, and the selection of subsequent actions [44, 45]. In PRISM-M, this prospective influence may therefore represent one functional feature of model-like organization. However, predictive information alone is not sufficient to define an internal neural model, because neural activity may contain decodable information without that information being functionally used [36, 37]. Stronger evidence would combine prospective information with integration, persistence or reinstantiation, and measurable effects on subsequent neural processing or behavior.

PRISM-M also intersects with predictive-processing theories, which propose that internal models generate predictions and that mismatches contribute to updating [9, 46]. The present framework uses prospective influence more broadly and does not require a specific prediction-error architecture. This distinction is consistent with recent analyses emphasizing that similar neural response patterns can arise from different computations and that apparent prediction-error signals require careful identification of the information and computation involved [47]. Prediction is one possible use of an internal model in PRISM, alongside simulation, inference, regulation, and action.

Comparison with two standard prediction models helped place the PRISM-M results in context. ARX provides a general system-identification framework in which future values are predicted from preceding states and current inputs [24], while a Kalman model estimates the evolution of an underlying state from noisy observations [23]. The ARX model performed almost as well as the full PRISM-M model, while the Kalman model was less accurate. This close correspondence is consistent with the ARX(1)-like recurrent structure of the present scalar realization. The contribution of PRISM-M lies in organizing this recurrent structure into biologically interpretable operations of extraction, compression, integration, stabilization, and subsequent model use. This decomposition provides explicit functional components that can be examined individually and tested experimentally as candidate processes involved in internal neural model formation. Future comparisons can therefore consider both predictive performance and the extent to which this structured organization helps explain neural computation and guide experimental testing [34].

While the simulations helped define how PRISM-M can generate persistence, contextual sensitivity, stable adaptation, and prospective information, the BART analysis provided an initial experimetal test of selected features of the framework using human fMRI data. Condition classification served as a basic check that the supplied spatial data retained task-related structure, while inflation-related activity showed a modest association with pumping behavior. Previous BART neuroimaging studies have identified distributed prefrontal, cingulate, insular, striatal, and related activity during sequential risk taking [9], providing an established neural context for the task. The present findings are compatible with structured, behaviorally relevant neural-state information. At the same time, the supplied fMRI files represented participant-level spatial features rather than temporally resolved trajectories, and static neural summaries did not reliably improve behavioral prediction beyond task-structure information. The data-based analysis therefore addressed selected spatial and behavioral features of the framework and helped define the temporally resolved measurements needed to test recurrent integration and state-to-state evolution.

Because the BART data consisted primarily of spatial neural summaries, the next step is to examine how PRISM-M states evolve over time. This is particularly important because the framework makes specific predictions about the temporal progression, stabilization, and updating of neural states. Contemporary approaches to neural dynamics similarly emphasize ordered trajectories, latent state spaces, and relationships between present and future population states [32–34]. Event-resolved fMRI could provide one approach when trial timing and hemodynamic modeling allow sequential state estimation. Electroencephalography (EEG) and magnetoencephalography (MEG) provide substantially greater temporal resolution, while intracranial electrophysiology, large-scale extracellular recordings, and calcium imaging can offer more direct measurements of evolving neural activity. Particularly informative studies would combine temporally resolved recordings with controlled changes in input and context and measurable downstream behavior. These studies could test a sequence of properties rather than relying on a single neural signature. A candidate state could first be evaluated for structured information about relevant signals, then for systematic contextual modulation, persistence or reconstructability across an appropriate interval, and contribution to a subsequent neural or behavioral state. Perturbation would add stronger functional evidence by asking whether altering the candidate state changes the predicted downstream consequence. This progression is consistent with current efforts to move from neural decoding toward functional and causal accounts of how representations are used [36, 37, 41]. Replication across tasks, cohorts, recording modalities, and neural systems will be important for determining which signatures generalize.

The present mathematical formulation was intentionally kept simple so that the relationships among the PRISM operations could be examined directly. Using scalar states and a limited number of parameters made it possible to test how extraction, compression, integration, and stabilization interact and to determine whether the resulting recurrent dynamics remain stable. The current PRISM-M model focused mainly on how the first four PRISM operations generate and maintain a model state, while Operation 5 was represented more simply by testing whether the established state contained information about the next state. Future studies can build on this framework by introducing multidimensional neural states, nonlinear interactions, multiple timescales, richer contextual inputs, and more complex forms of compression. Operation 5 can also be developed further to examine how an established state is reinstated, used for prediction or simulation, translated into action or regulation, and updated through feedback (Figure 6). Additional simulations can test how robust the findings are across broader parameter ranges and alternative model assumptions. A further step will be to estimate model parameters from experimental data and test whether PRISM-M can reproduce or predict temporal trajectories observed in neural recordings [34].

Overall, the present results suggest that a relatively simple recurrent model can generate several properties relevant to internal neural model formation. Represented input can be integrated with context, the resulting state can persist while remaining revisable, different balances between persistence and updating can emerge within recurrently stable dynamics, and the established state can carry information forward. These findings are consistent with concepts from recurrent-network theory, stability– plasticity, efficient and population coding, neural state-space approaches, predictive processing, and established internal-model frameworks, without requiring a single biological implementation. PRISM-M therefore provides a testable framework linking broad theories of representation and prediction with the neural dynamics through which internal models may be formed and used. Its value will depend on whether these relationships remain informative across richer computational models, independent datasets, temporally resolved neural recordings, and direct experimental manipulation.

## Data and Code Availability

The complete PRISM-M Version 4 reproducibility package, including source code, model specifications, simulation and analysis scripts, numerical source data, derived outputs, figure-source data, software requirements, provenance information, and SHA-256 manifests, is publicly available through Zenodo at DOI: **10.5281/zenodo.22070209**. The original BART behavioral and fMRI dataset analyzed in this study is available from Mendeley Data (Wei and Qin, 2025; DOI: 10.17632/xjn4n9cvxs.1).

